# Perinatal fluoxetine exposure increases male rat sexual behavior

**DOI:** 10.1101/2025.05.26.656113

**Authors:** Ane K. Baann, Danielle J. Houwing, Jocelien D.A. Olivier, Roy Heijkoop, Eelke M.S. Snoeren

**Affiliations:** Department of Psychology, UiT the Arctic University of Norway, Tromsø, Norway; Neurobiology, Groningen Institute for Evolutionary Life Sciences, University of Groningen, Groningen, the Netherlands

**Keywords:** fluoxetine, perinatal, sexual behavior, rats, seminatural environment

## Abstract

Selective serotonin reuptake inhibitors (SSRIs) are commonly prescribed to pregnant women due to their efficacy and safety profile, leading to potential exposure of the developing fetus and infant. Serotonin, which acts as a neurotransmitter in adults and is crucial for regulating male sexual behavior, also serves as a neurotrophic factor during early brain development. Yet, the long-term consequences of perinatal SSRI exposure on adult sexual functioning remain poorly understood. This study investigates the long-term effects of perinatal SSRI exposure on the sexual behavior of adult male rats in a seminatural environment. During pregnancy and lactating, mother rats were administered either fluoxetine (FLX, 10 mg/kg) or a control solution (CTR, 1% Methylcellulose) daily via oral gavage. Upon reaching adulthood, male offspring were assessed for sexual performance in a semi-natural setting, where they lived in mixed-sex groups for eight days. Comprehensive observations of sexual, social, and conflict behaviors were scoring from the first to the last copulatory behavior during the period in which female rats were sexually receptive. Our findings reveal that perinatal FLX exposure significantly increases sexual behavior in adult male rats, as evidenced by a higher total number of copulatory behaviors. This suggests that elevated serotonin levels during early development have enduring consequences for male rat sexual behavior in adulthood, potentially enhancing reproductive strategies and success in naturalistic environments.

## 1. Introduction

Selective serotonin reuptake inhibitors (SSRIs) are the most widely prescribed class of antidepressants in pregnant women due to their demonstrated efficacy, minimal adverse side effects, and favorable safety profile (Barbey and Roose, 1998; Gentile, 2015). Consequently, the prescription of SSRIs during pregnancy to support maternal mental health has become more prevalent, with approximately 2-7% of pregnant women in Canada and Western European countries using SSRIs (Charlton et al., 2015; Oberlander et al., 2006; Ververs et al., 2006; Zoega et al., 2015), and between 5-13% in Australia and the USA (Alwan et al., 2011; Andrade et al., 2008; Cooper et al., 2007). However, SSRIs have the capability to traverse the placental barrier and can be detected in breast milk (Kristensen et al., 1999; Rampono et al., 2004; Staal et al., 2024), thereby potentially affecting the developing fetus and infant (Heikkinen et al., 2003; Olivier et al., 2011).

By inhibiting the serotonin transporter, SSRIs prevent the reuptake of serotonin (5-HT) into presynaptic nerve terminals, leading to an elevation in synaptic 5-HT concentrations. In adults, 5-HT primarily functions as a modulatory neurotransmitter, influencing emotion, stress responses, sleep, learning, cognition, and attention (Canli and Lesch, 2007). Conversely, during early brain development, 5-HT also serves as a neurotrophic factor, regulating critical processes such as cell division, differentiation, migration, and synaptogenesis (Azmitia, 2001; Gaspar et al., 2003; Ramsteijn et al., 2022; Tate et al., 2021). Consequently, changes in 5-HT levels during in utero neurodevelopment are hypothesized to affect these developmental processes, similar to their impact on serotonergic function postnatally, potentially increasing susceptibility to affective disorders (Lesch and Mossner, 1998; Nadeem et al., 2022). Previous research in humans has shown changes in sleep patterns, social-emotional development, and internalizing and externalizing behavior in the offspring after perinatal SSRI exposure (Brandlistuen et al., 2015; Oberlander et al., 2010; Park et al., 2021; Weikum et al., 2013), as well as increased depressive and anxiety disorders into young adulthood (Bliddal et al., 2023; Rommel et al., 2021). Similarly, an increased risk for autism spectrum disorder was found in some studies (Andalib et al., 2017; Boukhris et al., 2016; Rai et al., 2013), although others did not find such a link or suggest that this increased risk is caused by the depression itself rather than the SSRIs (Brown et al., 2017; Hagberg et al., 2018; Hviid et al., 2013; Sorensen et al., 2013). Despite observed observations between SSRI exposure and child development, findings are complex and marked by inconsistent findings, raising the question whether these effects are due to the SSRIs or to the underlying maternal mental health condition (Suarez et al., 2022).

The benefit of conducting research with rodents is the ability to distinguish the effects of maternal SSRI use from the effects of maternal depression on the offspring. Extensive rodent research has been done to explore the effects of SSRI exposure on different social behaviors. As such, studies have delved into the impact of SSRI exposure during development on social play and communicating behavior in juvenile rats (Gemmel et al., 2017; Houwing et al., 2019b; Khatri et al., 2014; Olivier et al., 2011; Rodriguez-Porcel et al., 2011; Simpson et al., 2011), or social behavior (Gemmel et al., 2019; Houwing et al., 2019a; Houwing et al., 2019b; Ko et al., 2014; Olivier et al., 2011), prosocial behavior (Heinla et al., 2020), communication abnormality (ultrasonic vocalizations) (Houwing et al., 2019b; Lan et al., 2023), and aggressive behavior in adult rodents (Houwing et al., 2020; Kiryanova and Dyck, 2014; Svirsky et al., 2016).

Furthermore, studies have shown that perinatal SSRI exposure can negatively affect the motivation to start social contact (Khatri et al., 2014; Rodriguez-Porcel et al., 2011; Simpson et al., 2011; Zimmerberg and Germeyan, 2015), but not novelty-induced social investigation behaviors (Sylte et al., 2021).

The impact of perinatal SSRI exposure on male sexual behavior has not been thoroughly investigated in rats. However, some studies using postnatal SSRI exposure have shown that elevated 5-HT levels during development can inhibit male rat sexual behaviors at adulthood (Harris et al., 2012; Rayen et al., 2013; Rodriguez-Porcel et al., 2011). Previous studies in our labs, however, showed both that perinatal SSRI exposure has no effect (Houwing et al., 2020) and that it leads to increased mount and intromission activity in males (Houwing et al., 2019a), while no effects were found on sexual behavior of female rats (Hegstad et al., 2020; Houwing et al., 2019a).

A limitation of most of the previous studies, however, is the use of traditional test set-ups in which male and female rats have limited amounts of space and time to interact with each other. In addition, testing rats in dyads prevents us from exploring social interaction in a more natural setting where rats live in complex social groups (Calhoun, 1963; Robitaille and Bovet, 1976; Schweinfurth, 2020), usually in a burrow system. In nature, a colony of rats consists of one or several females and males. Interestingly, the mating patterns in rat groups differ significantly from those seen in traditional lab tests with rat pairs. In group settings, multiple males and females mate simultaneously (Chu and Ågmo, 2014, 2015b; McClintock, 1984; McClintock and Adler, 1978; Snoeren et al., 2015). To understand the effects of perinatal SSRIs on sexual behavioral outcomes in a more natural setting, a semi-natural environmental setup was used. This allowed rats to exhibit their full range of natural behaviors and enabled us to investigate sexual behavioral changes within a natural social context. (Bove et al., 2018; Buwalda et al., 2017; Le Moene and Agmo, 2018; Le Moëne and Ågmo, 2019).

In our previous research, we demonstrated that male rats exposed perinatally to fluoxetine exhibit increased mounts and intromissions in a seminatural environment compared to control rats (Houwing et al., 2019a). However, this study was limited to observing only a random 30- minute segment of their sexual activity. Inspired by the Chu and Ågmo studies, who showed that females and males copulate for a longer period (Chu and Ågmo, 2014, 2015b), and that male rats copulate in multiple copulation bouts as long as receptive females remain available (Chu and Ågmo, 2015b), we re-analyzed the data of our previous study (Houwing et al., 2019a).

Based on the fact that female rats appear to have a longer behavioral estrus that is subdivided into multiple copulation bouts (Hegstad et al., 2020), the aim of this study was to re-analyze the data of the entire copulatory phase of male rats, measured from the first copulatory behavior until the last, and to determine the effects of perinatal fluoxetine exposure of male rat sexual behavior.

## 2. Material and Methods

The data presented in this article consists of a more detailed analysis of a previous study (Houwing et al., 2019a). The methods in this manuscript are similar to the other study, but adapted to specifically fit the aim of this study.

### 2.1 Animals and dam housing conditions

Upon arrival, the rats (Charles River in Sulzfeld, Germany, weighing 200-250 g at the time of arrival) were allowed to acclimatize to the facility for one week, before breeding was started. Breeding was conducted with ten male and ten female Wistar rats. Unless stated otherwise, all rats were housed in same-sex pairs in Makrolon©IV cages with a 12:12 h light/dark cycle (lights on at 11:00 AM), a room temperature of 21 ± 1°C, and humidity levels of 55 ± 10%. Standard chow (Special Diet Service) and tap water was provided ad libitum. Nesting materials and (other) environmental enrichment were also provided. All experimentation was carried out in agreement with the European Union council directive 2010/63/EU. The protocol was approved by the National Animal Research Authority in Norway (project 8314) on the 16^th^ of March 2016.

### 2.2 Breeding and antidepressant treatment

On the day of proestrus, a female and male rat were housed together for 24-hours. The start of proestrus was assessed by placing them together and confirming the display of lordosis behavior. After the 24-hour mating period, both male and female rats returned to their original home cages with a same-sex conspecific. Throughout gestational days 1 to 14, the pregnant dams were housed with a same-sex partner. From gestational day 14, the pregnant females were relocated to individual housing in Makrolon©IV cages with access to nesting materials, and twice-daily assessments were performed to monitor pup delivery for each mother. After birth, pups remained housed with their mothers until weaning on postnatal day 21 (PD21).

From gestational day 1 (G1) to postnatal day 21 (PND21), dams received daily oral gavage treatment with either 10mg/kg fluoxetine (FLX, Apoteksproduksjon, Oslo, Norway) or a control (CTR, 1% Methylcellulose, Sigma, St. Louis, MO, USA) with a stainless-steel feeding tube. The fluoxetine treatment, was prepared by dissolving pulverized tablets made for human consumption, resulting in a 2 mg/mL concentration and administration at 5 mL/kg.

Methylcellulose powder, a non-active filler, was dissolved in water to create a 1% solution for the vehicle, also administered at 5 mL/kg. Regular weight assessments occurred every third day, with adjustments to dosage as needed. The fluoxetine dose was determined based on literature comparing human and animal blood levels(Lundmark et al., 2001; Olivier et al., 2011).

### 2.3 Offspring housing conditions before the seminatural environment

On PND21, the offspring were weaned, ear punched for individual identification and housed in pairs or trios of same-sex littermates in Makrolon IV cages. Until an age of 13-18 weeks, the offspring was left alone with minimal interference, limited to weekly cage cleaning. Only the female offspring were ovariectomized two weeks before the start of the experiment, after which they were single housed for 3 days before returning to their home cage.

### 2.4. Ovariectomy surgery

Ovariectomy was performed on the female offspring so that we were able to control of the onset of behavioral receptivity with hormone injections. The ovariectomy was performed under isoflurane anesthesia and analgesia with buprenorphine (0.05 mg/kg, subcutaneously) and Carprofen (5 mg/kg, subcutaneously). A 1–2 cm longitudinal midline skin incision was made at the lower back of the female, followed by bilateral muscle incisions above the ovaries. After that the ovaries were extirpated and the fallopian tube stump was ligated. The muscle incisions were sutured and a wound clip was placed for skin closure. The females received post-surgery analgesia with 5 mg/kg Carprofen (subcutaneously) 24 and 48 h after surgery and were monitored daily for a week.

### 2.5 Seminatural environment

The seminatural environment (2.4 x 2.1 x 0.75 m) consisted of an interconnected burrow system and an open field area linked by four 8 x 8 cm openings. A visual representation can be found in (Houwing et al., 2019a; Snoeren et al., 2015; Sylte et al., 2021). The burrow system had tunnels (7.6 cm wide, 8 cm high), four nest boxes (20 x 20 x 20 cm), and was covered with Plexiglas.

The open area featured partitions (40 x 75 cm) simulating natural obstacles. A curtain between the open field and burrow area allowed separate light control with complete darkness in the burrow and a simulated day-night cycle in the open area. A lamp (2.5 m above the center) emitted light (180 lx) from 10:45 pm to 10:30 am, simulating daylight. Flooring in both areas had a 2 cm layer of aspen wood chip bedding. Nest boxes included nesting material, shelters, and aspen wooden sticks, and food and water were available *ad libitum* in front of the open area wall (2 kg of pellets were provided at the start of the experiment). Video cameras (Balser), positioned 2 m above the arena’s, recorded 24-hour cycles.

### 2.6 Procedure

The day before their introduction to the seminatural environment, offspring were shaved and marked with tail marks under isoflurane anesthesia for individual recognition on video. At 10 a.m. on the first day of the experiment (day 0), a cohort of 4 male and 4 female offspring (from different litters and homecages) was placed in the seminatural environment for 8 days (Figure 1). Notably, all rats entered the seminatural environment in a sexually inexperienced state. Each cohort was composed of 2 control (CTR)-females, 2 intact CTR-males, 2 fluoxetine (FLX)- females and intact 2 FLX-males. In total, five cohorts of rats were introduced into the seminatural environment, resulting in treatment groups of n=10.

**Figure 1.**
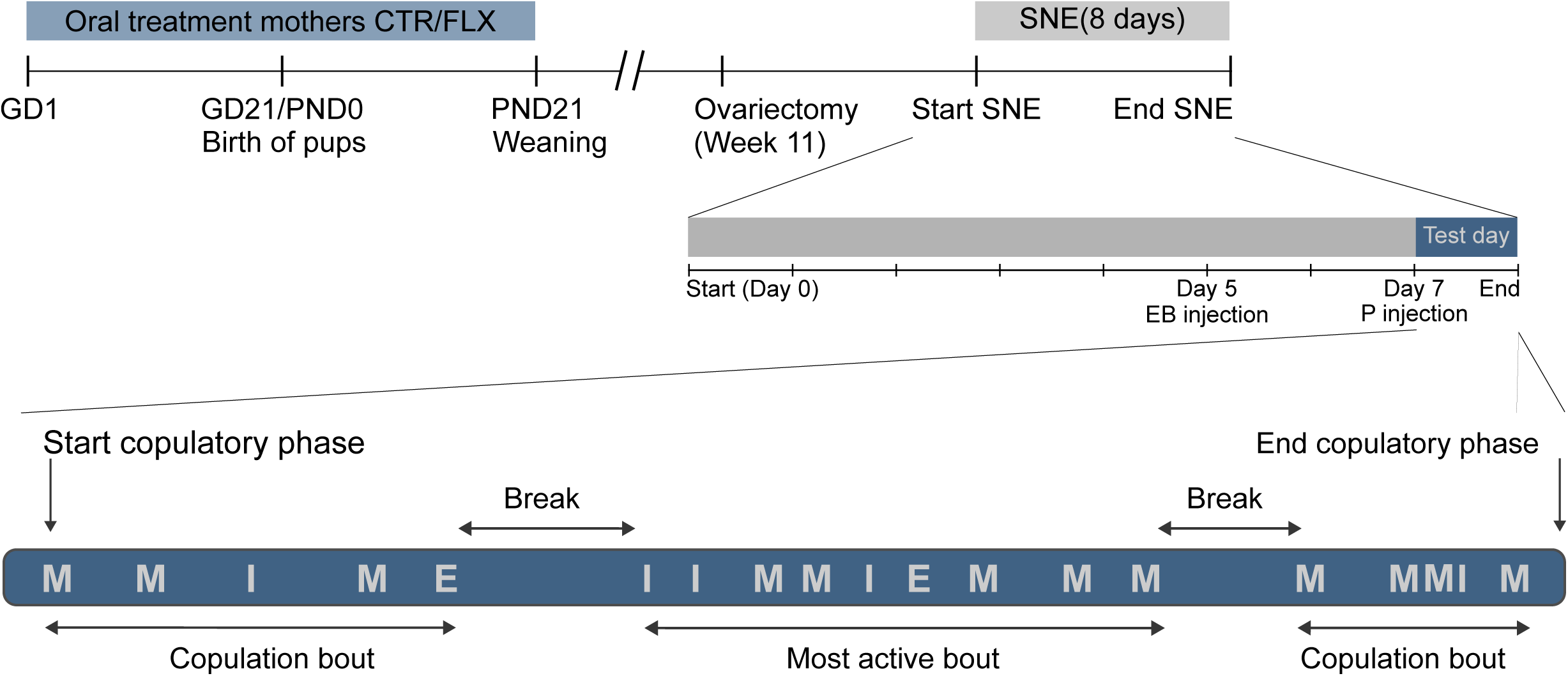
Schematic overview of the design of the experiment. Mothers were orally administered with either 10 mg/kg fluoxetine (FLX)or control (CTR) treatment from gestational day (GD) 1 until weaning on postnatal day (PND) 21. The offspring were left undisturbed until adulthood. Female offspring were ovariectomized at week 11 At 13-18 weeks of age, both males and females were placed in the seminatural environment (SNE) for 8 days in groups of 8 rats (2 FLX females, 2 CTR females, 2 FLX males, 2 CTR males). Male sexual behavior was scored at day 7, after the females received estradiol benzoate (EB) and progesterone (P). An example of what a copulatory phase could look like, with copulation bouts and breaks consisting of mounts (M), intromissions (I) and ejaculations (E). The most active bout is the bout in which most copulatory behaviors were performed.

To synchronize receptivity among all female rats on the same day of observation in the seminatural environment, estradiol benzoate was administered to females on day 5 at 10:00 am, followed by progesterone on day 7 at 10:00 am. Subsequently, at 10 am on day 8, the experiment ended, and the rats were taken out of the seminatural environment. The environment was carefully cleaned and prepared for the next cohort of rats.

### 2.7 Behavioral analysis

A trained observer reviewed all video recordings of day 7 and day 8 to identify the initial display of copulatory behavior (mount, intromission, or ejaculation) of each male during the receptive phase of the female rats (that can last 24 hours). The 1^st^ copulatory behavior marked the start of the copulatory phase (Figure 1). Like the previous study on the female rats(Hegstad et al., 2020), a copulatory phase consisted of different copulation bouts. A bout consisted of 1 or more mounts, intromissions and ejaculations, and started from the first and ended with the last copulatory behavior, after which a break of at least one hour of copulatory-related inactivity took place (Figure 1).

A trained observer annotated the start and end of a copulation bouts, mounts, intromissions, and ejaculations. Moreover, social (sniffing others) and conflict (nose-off, boxing/wrestling, and flighting) behaviors were also scored during each copulation bout (for more details, see(Hegstad et al., 2020)). Although not further mentioned in this manuscript, we also annotated the behavior of the partner with which these behaviors took place and the location of the behavior (burrow, open field). The annotations were performed with Observer XT, version 16 (Noldus, Wageningen, the Netherlands). The data was analyzed with a customized python script.

Unfortunately, a (blue) marker not visible under fluorescent light was used in one of the cohorts. This made it difficult to recognize individuals. Therefore, we only observed the copulatory behaviors in this cohort and did not analyze social and conflict behaviors. This resulted in a total sample size of ten CTR- and ten FLX-males for copulatory analysis and eight CTR- and eight FLX-males for social and conflict behavioral analysis.

### 2.8 Data preparation and statistical analysis

First, we analyzed the data of the entire *copulatory phase* that consisted of the data of all *copulation bouts* together, without the breaks in between them. Then, we determined the *most active copulation bout* for each rat as the copulation bout in which most copulatory behaviors were performed. The duration, total number of copulatory behaviors and time spent on social and conflict behaviors were calculated for both the entire copulatory phase and the most active copulation bout.

For the most active copulation bout for each male, we also explored various measures of copulatory efficiency. The *intromission ratio* was calculated by dividing the number of intromissions by the total number of intromissions and mounts. The *inter-intromission interval* was calculated by dividing the number of intromissions by the copulation bout duration. The *copulatory rate* was calculated by summarizing the number of mounts and intromissions and dividing it by the duration of the bout. Finally, the *latency to ejaculate* was computed for the most active copulation bout by tracking the time of the first mount or intromission to the time to the first ejaculation. We also calculated the latency to the 1^st^ ejaculation of the rat (of the entire copulatory phase), by calculating the ejaculation latency of the copulation bout that contained the 1^st^ ejaculation and add the duration of preceding copulation bouts if this wasn’t the 1^st^ copulation bout. It should be mentioned that these parameters could only be calculated if the rat had at least 1 ejaculation. The rats without ejaculations were excluded from these parameters.

The homogeneity of variance was tested with the Shapiro–Wilk test. The FLX males were compared to CTR males with an independent *t-*test if the data was normally distributed, whereas the non-parametric Mann–Whitney U test was used for the other non-normally distributed data.

In order to analyze the patterns of sociosexual behavior over the full course of the copulatory phase and the most active copulation bout, we divided the total duration into 5%-time bins. Then we analyzed the behaviors performed in each 5%-time bin over the full course of the phase and bout. A mixed models analysis with Treatment (CTR vs FLX) and Time (5%, 10%, 15%, etc. to 100%) as factors was performed, followed Bonferroni corrected post-hoc t-tests in case of significance (p<0.05).

## 3. Results

### 3.1 Sexual behavior

As mentioned in the methods, the copulatory phase of the male rats was divided in multiple copulation bouts, with the copulation bout including most copulatory behaviors being defined as the ‘most active bout’. As shown in Figure 2A, FLX males started mating around the same time as CTR males (Z=-0.567, p=0.574), and there was no difference in the start of the ‘most active bout’ (Fig. 2B, Z=-0.529, p=0.597However, the most active bout of FLX males was significantly longer than that of CTR males (Fig. 2C, Z=-2.420, p=0.016). In addition, they mounted (Fig. 2D, Z=-2.571, p=0.010), intromitted (Fig.2E, Z=-2.540, p=0.011), and ejaculated (Fig. 2F, Z=-2.026, p=0.043) more often than CTR males in the entire copulatory phase. No difference in the latency to 1^st^ ejaculation (calculated from the start of the 1^st^ copulation bout until the 1^st^ ejaculation, excluding the time of the breaks in-between bouts) between FLX and CTR males were found (*t*=-2.076, p=0.072, data not shown).

**Figure 2.**
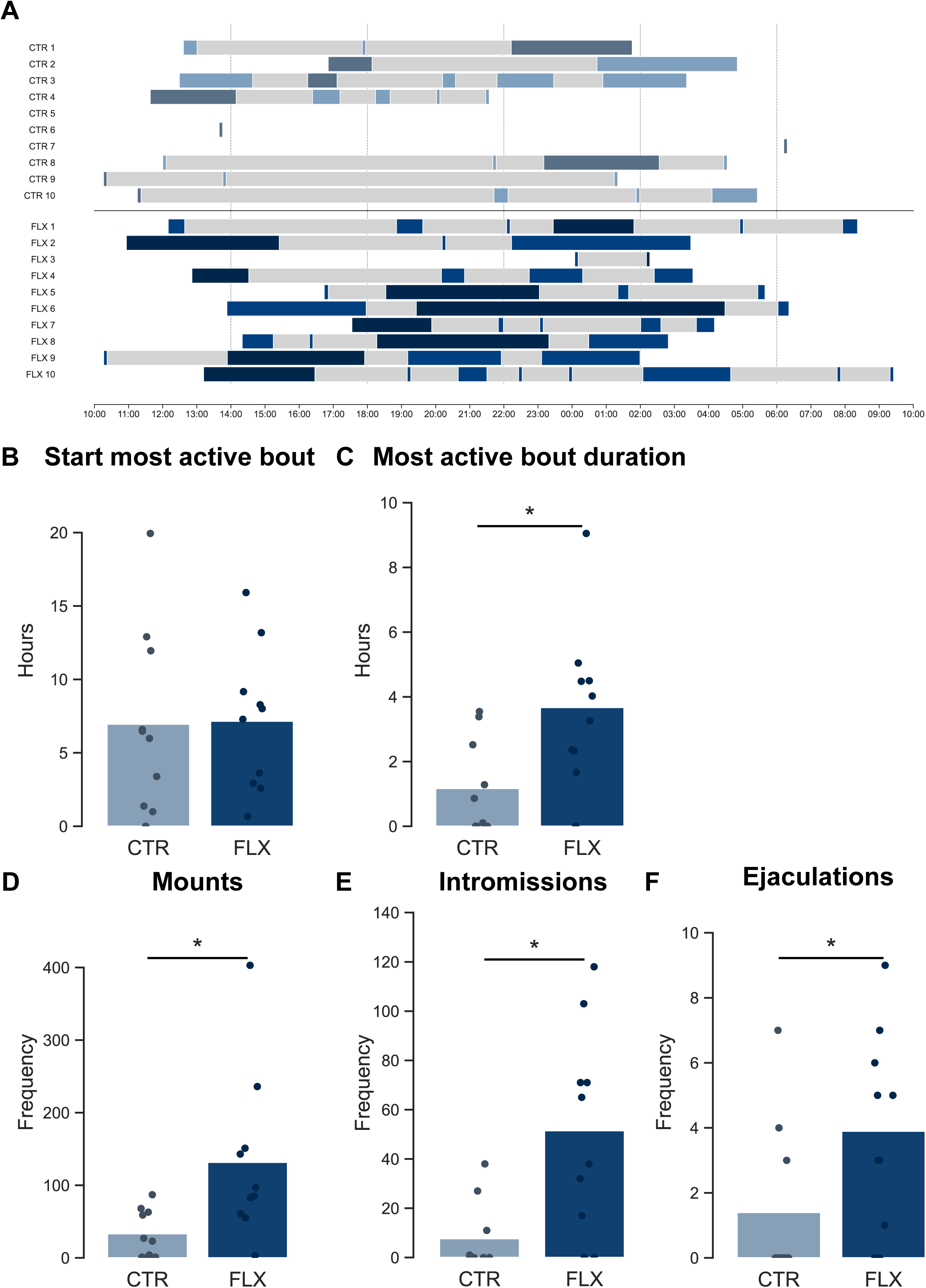
Effects of perinatal fluoxetine exposure on the copulatory phase of male rats. A) Schematic overview of the timing, duration and number of different copulation bouts. The intermitting periods of sexual inactivity are shown in light grey and the copulation bouts in light blue (CTR-males, upper part) or dark blue (FLX-males, lower part). The most active bouts are shown in darker colors. The copulatory phase starts with the first mount, intromission or ejaculation and thus with the onset of the first copulation bout, and ends with the last mount, intromission or ejaculation and thus at the end of the last copulation bout, B) the onset of the most active bout in seconds after progesterone injection (at 10:00 am), C) the duration of the “most active bout” in seconds. D) number of mounts in the total copulatory phase, E) number of intromissions in the total copulatory phase, F) number of ejaculations in the total copulatory phase. Data are shown with individual data points, with the bars representing the mean.

By looking at the ‘most active bout’ in more detail, we are able to analyze more specific copulatory parameters, such as the inter-intromission interval, copulation rate, intromission ratio and latency to 1^st^ ejaculation. Just as in the entire copulatory phase, FLX males mounted (Fig. 3A, Z=-2.535, p=0.011) and intromitted (Fig.3B, Z=-2.378, p=0.017) more often than CTR males across the entire ‘most active bout’. With regard to ejaculation, however, no difference between FLX and CTR males was observed (Fig. 3C, Z=-1.583, p=0.143), meaning that they needed more mounts and intromissions to reach an ejaculation. Interestingly, the latency to achieve the 1^st^ ejaculation within this most active bout (measured from the 1^st^ copulatory behavior) was longer in FLX than CTR males (Fig. 3D, Z=-2.393, p=0.017). No differences were found in the inter-intromission intervals (Fig. 3E, *t*=0.453, p=0.661), intromission ratio (Fig. 3F, Z=-1.612, p=0.107) or copulation rate (Fig. 3G, Z=-1.612, p=0.650) calculated for the total ‘most active bout’. When we calculate these parameters until the 1^st^ ejaculation in the ‘most active bout’, we found again no differences in intromission ratio (Z=-1.527, p=0.127, data not shown) or copulation rate (Z=-1.134, p=0.257, data not shown). Now, however, FLX males had longer inter-intromission intervals during the 1^st^ ejaculatory series of the ‘most active bout’ than CTR males (Fig. 3G, *t=*-2.582, p=0.030).

**Figure 3.**
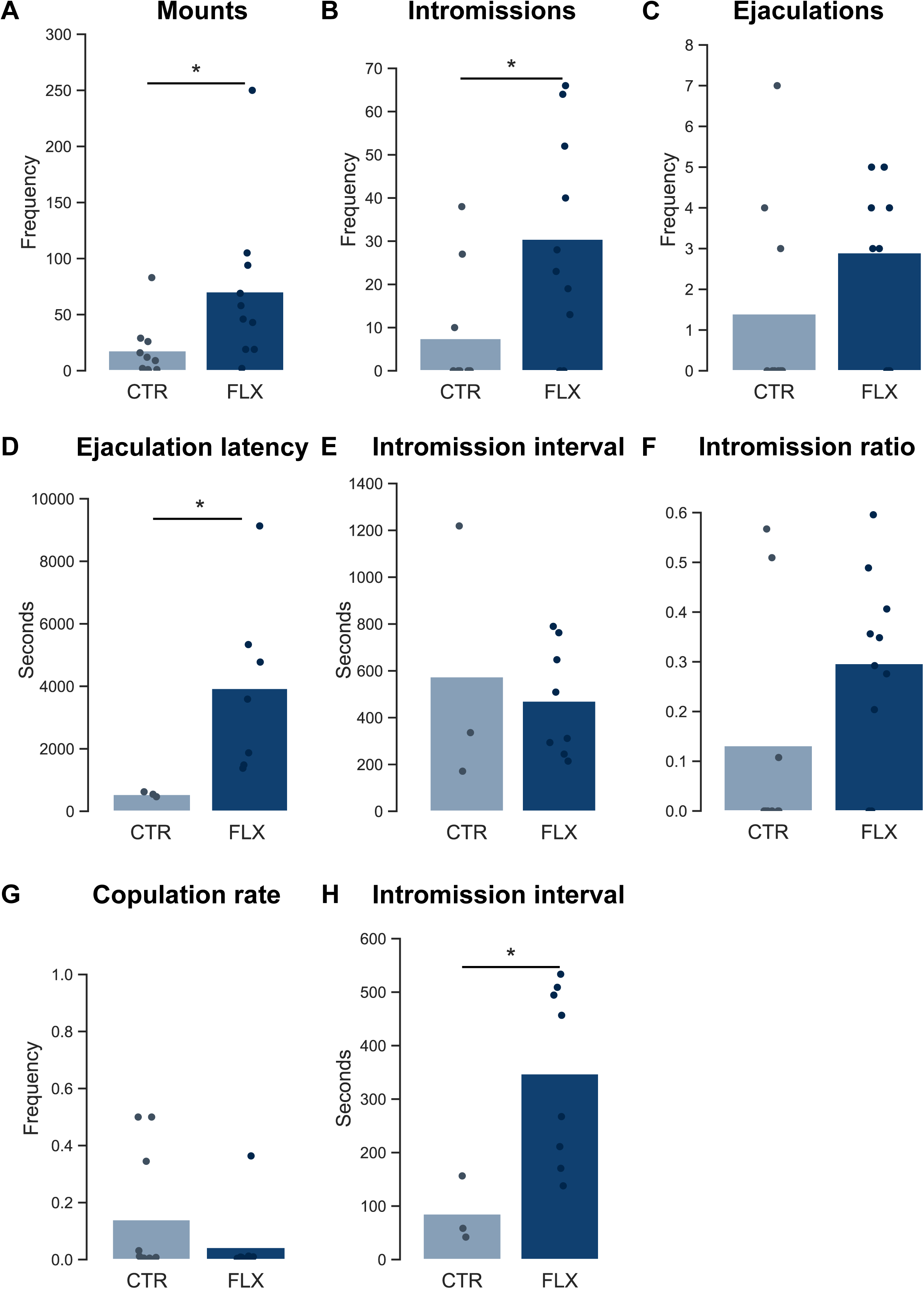
Effects of perinatal fluoxetine exposure across the most active bout of male rats. A) number of mounts, B) number of intromissions, C) number of ejaculations, D) latency to 1^st^ ejaculation within the bout, measured from the 1^st^ mount or intromission of the bout E) intromission interval for complete bout, F) intromission ratio, G) copulation rate, H) intromission interval for the 1^st^ ejaculatory series only. Data are shown with individual data points, with the bars representing the mean.

### 3.2 Social and conflict behavior

To gain deeper insights into the temporal dynamics of copulatory behavior, we analyzed the distribution of copulations throughout the entire copulatory phase and the most active bout. To facilitate comparison between individual animals, the data was segmented into 5% time bins. As illustrated in Figure 4, FLX rats exhibited a constant higher frequency of copulatory behavior compared to CTR males over the duration of the copulatory phase (Fig 4A, Treatment: F_(1,18)_=8.707, p=0.009, Time: F_(19,342)_=2.336, p=0.001, Treatment * Time: F_(19,342)_=0.827, p=0.674) and most active bout (Fig 4B, Treatment: F_(1,18)_=6.432, p=0.021, Time: F_(19,342)_=1.127, p=0.322, Treatment * Time: F_(19,342)_=1.202, p=0.253).

**Figure 4.**
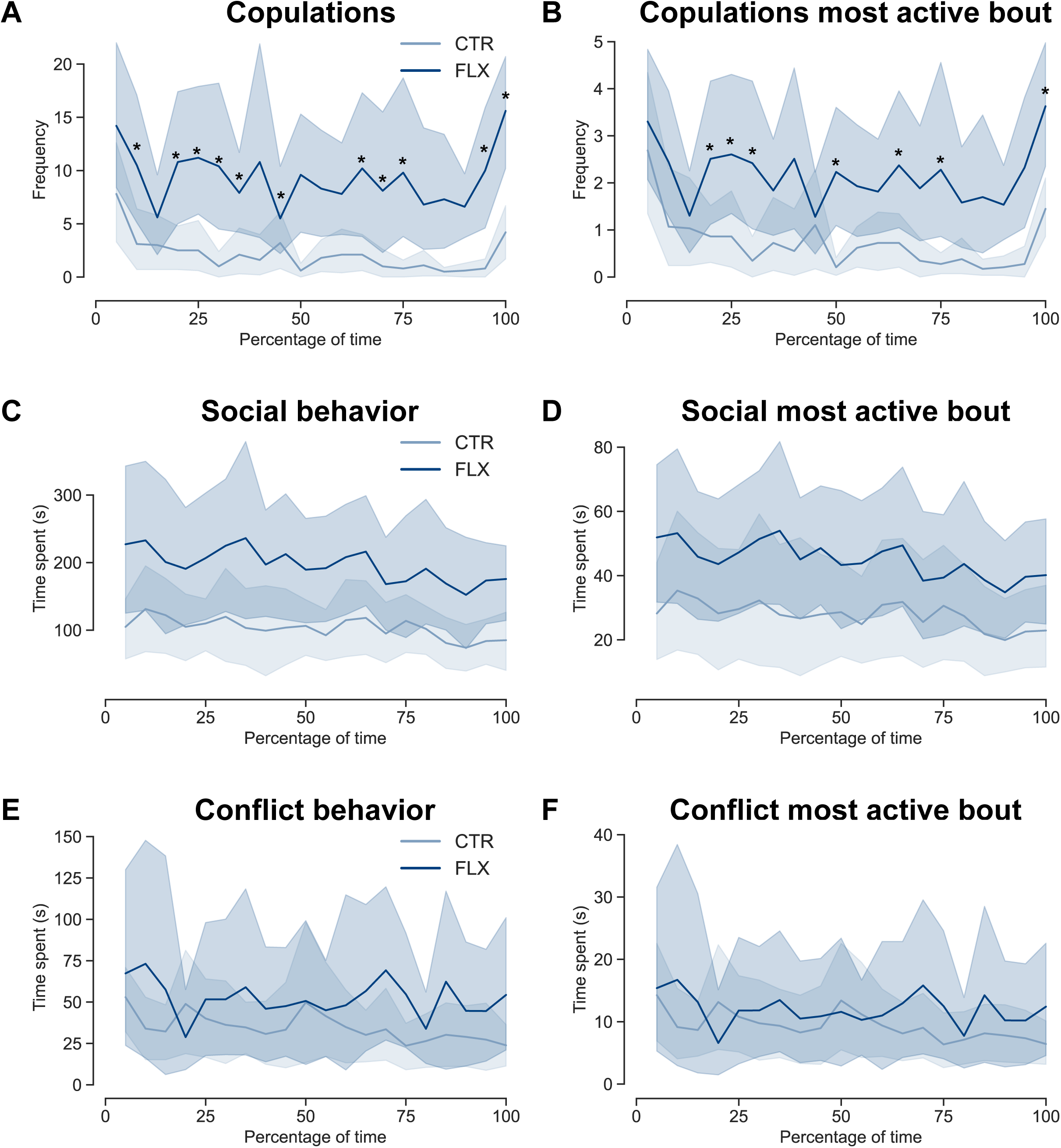
Effects of perinatal fluoxetine exposure on copulations, social and conflict behaviors over the course of the total copulatory phase and most active bout of male rats. A) number of copulations in total copulatory phase, B) number of copulations in most active bout, C) time spent on social behaviors (sniffing others) in total copulatory phase, D) time spent on social behavior in most active bout, E) time spent on conflict behaviors in total copulatory phase, F) time spent on conflict behaviors in most active bout. Data are calculated and visualized per 5% time bin of the total copulatory phase (left graphs) or most active bout (right graphs). The data are shown as mean (line) ± sem (shadow) for CTR- (light blue) and FLX-males (dark blue). * p<0.05 compared to CTR males of that time bin.

To confirm that the differences in sexual behavior were not caused by differences in social ability, we measured the time spent sniffing other rats (males and females) and showing conflict-related behavior (e.g. fighting, boxing and wrestling, nose-off). No differences in time spent sniffing others were found between FLX and CTR males over the course of the entire copulatory phase (Fig 4C, Treatment: F_(1,13)_=3.120, p=0.101, Time: F_(19,247)_=1.786, p=0.025, Treatment * Time: F_(19,247)_=0.378, p=0.992, no significance in post-hoc analysis for Time), nor in the most active bout (Fig 4D, Treatment: F_(1,13)_=1.967, p=0.184, Time: F_(19,247)_=1.425, p=0.115, Treatment * Time: F_(19,247)_=0.648, p=0.866). Similarly, FLX males spent an equal amount of time in conflict behavior over the course of the copulatory phase (Fig 4E, Treatment: F_(1,13)_=0.550, p=0.471, Time: F_(19,247)_=0.647, p=0.868, Treatment * Time: F_(19,247)_=0.657, p=0.859), or the most active bout (Fig 4F, Treatment: F_(1,13)_=1.847, p=0.197, Time: F_(19,247)_=0.450, p=0.978, Treatment * Time: F_(19,247)_=0.484, p=0.967).

## 4. Discussion

The main aim of this study was to conduct a comprehensive analysis of the effects of perinatal fluoxetine (FLX) exposure on sexual behavior of male rats under semi-natural conditions. We focused on the entire copulatory phase, which begins with the first display of copulatory behavior (mount, intromission, or ejaculation) and concludes with the last copulatory behavior (mount, intromission, or ejaculation) after hormone priming of the females. Our analysis revealed that both control (CTR) and FLX male rats copulate in multiple copulatory bouts, interspersed with breaks lasting at least one hour, during which no copulatory behaviors occur. Overall, we found that FLX-exposed males exhibited a higher frequency of mounts, intromissions, and ejaculations compared to CTR males, despite both groups having similar latencies to achieve their first ejaculation. This increased copulatory activity in FLX males was also evident during the most active bout, which contained the highest number of copulatory behaviors. Although both CTR and FLX males ejaculated a similar number of times during this most active bout, FLX males initially demonstrated a longer ejaculation latency, accompanied by longer intromission intervals during this 1^st^ ejaculatory series. However, once an ejaculation was achieved, no further differences in copulatory efficiency metrics, such as copulation rate, intromission interval, and intromission ratio, were observed between the two groups over the course of the most active bout. The elevated levels of copulatory behavior in FLX males remained consistent throughout the entire copulatory phase and the most active bout, and were not caused by disturbances in social or conflict behaviors.

These findings align with our previous observations in the same cohort of rats, where we analyzed 30 minutes of their behavior (Houwing et al., 2019a). However, our results contrast with other studies that reported reduced mounts, intromissions, and ejaculations following FLX exposure during development (Gouvea et al., 2008; Harris et al., 2012; Rayen et al., 2013; Rodriguez-Porcel et al., 2011), or that found no effect on male copulatory behavior (Cagiano et al., 2008; Olivier et al., 2011). This discrepancy could be attributed to the timing of FLX exposure or the test settings. Inhibitory effects were observed in studies in which FLX was administered postnatally (Gouvea et al., 2008; Harris et al., 2012; Rayen et al., 2013; Rodriguez-Porcel et al., 2011), while no effects were reported in prenatal exposure studies (Cagiano et al., 2008; Olivier et al., 2011). In contrast, the males in our study were exposed to FLX perinatally, which may account for the differences in outcomes. Alternatively, the differences could also be ascribed to the short test durations used in other studies. In our investigation, FLX-exposed males had a longer ejaculation latency during the most active bout. If the observation period had been terminated after the first ejaculatory series, the outcomes might have been interpreted differently. The time to reach ejaculation is longer, and therefore perinatal fluoxetine exposure might have been interpreted as making male rats less sensitive and therefore needing more mounts and intromissions. However, even though male rat sexual behavior seems less efficient to reach an ejaculation in this phase of the most active bout, the overall effect of perinatal fluoxetine exposure seems to boost male rat sexual behavior.

Our study benefited from the use of a seminatural environment, which allowed rats to exhibit the full range of their natural behaviors. The mating patterns observed in groups of rats differ significantly from those seen in traditional laboratory mating tests, which typically involve pairs of rats or mice (Chu and Ågmo, 2014, 2015b; Garey et al., 2002; McClintock, 1984; McClintock and Adler, 1978; McClintock and Anisko, 1982; McClintock et al., 1982; Snoeren et al., 2015). The use of the seminatural environment enabled us to assess the effects of perinatal SSRI exposure on natural mating behavior throughout the entire copulatory phase. This approach contrasts with the short copulation tests commonly used in previous studies. Additionally, this approach revealed that both CTR- and FLX- males displays sexual behaviors in multiple copulation bouts. This finding aligns with the comprehensive studies conducted by other colleagues, who examined the full range of sexual behavior patterns in male and female rats within a seminatural environment, using both intact and ovariectomized Wistar females (Chu and Ågmo, 2014, 2015a, b; Le Moëne et al., 2020).

The observed increase in sexual behavior among male rats perinatally exposed to FLX in a semi-natural setting raises important questions about the functional implications of such behavioral changes. In a natural environment, where competition for mates can be intense, the ability to engage in more frequent sexual behaviors could theoretically enhance reproductive success. Although FLX-exposed rats exhibited reduced copulatory efficiency and an increased delay in achieving their first ejaculation during the most active bout, they performed more mounts and intromissions and ultimately achieved more ejaculations than the control males throughout the entire copulatory phase. Given that multiple females are present in nature and also engage in multiple copulation bouts, receiving several ejaculations (Chu and Ågmo, 2014; Hegstad et al., 2020), a male capable of achieving more ejaculations would theoretically have a higher likelihood of reproductive success. Therefore, our findings indicate that perinatal exposure to SSRIs has the potential to enhance male reproductive strategies and success in naturalistic settings. However, this will ultimately depend on whether or not FLX exposure affects sperm quality, which needs to be further investigated in future research.

For the current study the purpose of observing other social behaviors was to assure that potential differences in sexual behavior were not attributable to abnormalities in social or conflict behavior. Since no significant differences were found between CTR- and FLX-males, we can conclude that the stimulatory effects of perinatal FLX exposure on sexual behavior were not influenced by potential effects on social (sniffing) or conflict behaviors. This aligns with our previous findings, which also showed no effect of FLX exposure on active social or conflict behavior in these males (Houwing et al., 2019a). However, other studies have reported conflicting results regarding the impact of developmental FLX exposure on social behavior.

Some studies have found that FLX exposure can increase social interactions (Ko et al., 2014), while others have reported decreased time spent on social exploration behavior (Houwing et al., 2019b; Olivier et al., 2011). A plausible explanation for these discrepancies is that our current experiment observed social behavior during the behavioral estrus phase of the females. During this period, sexual interaction becomes the primary focus of the rats, which could overshadow the effects of perinatal FLX exposure on other behaviors, such as social and conflict behavior. Moreover, some discrepancies in copulatory behaviors might be found by testing rats as virgin (Rayen, Steinbusch et al. 2013), or as sexually experienced rats (Houwing et al.,2019).

## 5. Conclusion

In conclusion, our study demonstrates that perinatal FLX exposure significantly enhances sexual behavior in male rats under semi-natural conditions, as demonstrated by increased frequencies of mounts, intromissions, and ejaculations compared to controls. Despite an initial delay in achieving the first ejaculation, FLX-exposed males exhibited increased overall copulatory activity without disturbances in social or conflict behaviors. These findings suggest that perinatal SSRI exposure may enhance male reproductive strategies and success in naturalistic environments. The use of a semi-natural setting allowed for a comprehensive assessment of natural mating behaviors, highlighting the potential for perinatal FLX exposure to positively influence male reproductive fitness.

## Acknowledgements

Financial support was received from Helse Nord #PFP1295-16, Norway. DJH was supported by the KNAW ter Meulen travel grant, the Netherlands. We also would like to thank Ragnhild Osnes, Carina Sørensen, Nina Løvhaug, Katrine Harjo, and Remi Osnes for their excellent caretaking of the animals.

